# Integrated Ultrasound Neuromodulation and Optical Neuroimaging in Awake Mice using a Transparent Ultrasound Transducer Cranial Window

**DOI:** 10.1101/2025.02.19.638722

**Authors:** Shubham Mirg, Krishnendu Samanta, Haoyang Chen, Jin Jiang, Kevin L. Turner, Fatemeh Salehi, Kathiravan M. Ramiah, Patrick J. Drew, Sri-Rajasekhar Kothapalli

## Abstract

Ultrasound neuromodulation is a rapidly advancing, non-invasive technique with significant therapeutic potential for treating various neurological disorders. Although extensive *in vitro* and *in vivo* studies have provided valuable insights into its modulatory effects, the underlying mechanisms remain poorly understood, limiting its clinical translation. Optical neuroimaging techniques can help investigate these mechanisms; however, the opacity and bulkiness of conventional ultrasound transducers pose significant challenges for their integration with *in vivo* ultrasound neuromodulation studies, particularly in awake rodents. To address these limitations, we propose a straightforward solution: a miniaturized lithium niobate-based transparent ultrasound transducer (TUT) integrated as a thinned-skull cranial window for ultrasound stimulation while facilitating multimodal optical neuroimaging in awake mice brain. Using laser speckle contrast imaging and intrinsic optical signal imaging, we studied changes in brain hemodynamics in response to various ultrasound stimulation sequences. Our experiments demonstrated that TUT cranial window can robustly induce neuromodulatory effects with observed increase in both cerebral blood flow and total hemoglobin, with peak and cumulative hemodynamic changes directionally correlated with ultrasound stimulation duration and intensity. Overall, these findings highlight that TUT cranial window can seamlessly integrate ultrasound stimulation and optical neuroimaging in awake mouse brain models, offering promising prospects for uncovering the underlying mechanisms of ultrasound neuromodulation.

## 1 Introduction

The ability to stimulate brain and regulate its functions is key to understanding various neurological processes and treating neurological disorders. Stimulation techniques based on optical, electrical, and magnetic fields have been explored to modulate neural activity at varying depths and spatial resolutions [1], [2]. More recently, ultrasound stimulation has garnered significant attention due to its ability to target deep brain regions non-invasively and has been studied as a therapeutic intervention for various diseases, including epilepsy, depression, Alzheimer’s disease [3], [4], [5]. Studies have shown that ultrasound stimulation modulates neuronal activity [6], [7], [8] and induces hemodynamic changes via neurovascular coupling [9]. However, a comprehensive understanding of its mechanisms remains lacking, thereby hindering its rapid adoption and clinical translation.

Techniques such as magnetic resonance imaging, positron emission tomography and functional ultrasound imaging have been used to study ultrasound neuromodulation [10], [11], [12], [13], but they are limited in spatial (<10 μ m) and temporal resolutions (<100 ms) thereby hindering investigation of molecular mechanisms. Optical imaging, on the other hand, with its multiple endogenous and exogenous chromophores, is particularly well suited for studying mechanism of ultrasound neuromodulation. However, since the ultrasound stimulus is generated by an opaque and bulky transducer that requires an acoustic-friendly coupling medium (e.g., water or ultrasonic gel), various technical challenges and considerations arise when integrating optical imaging with conventional ultrasound transducers. Often, oblique geometries are necessary, which can lead to non-uniform stimulation, difficulties in precise targeting, and a lack of space at the imaging head—factors that ultimately limit multimodal integration [6], [7], [14]. Additionally, the use of water or gel as a coupling medium can introduce temperature confounds by acting as a heat sink, causing fluctuations in the temperature around the brain region [15].

The large and bulky nature of ultrasound neuromodulation setups further poses significant challenges for researchers in designing rodent studies. In some cases, the rodents undergo craniotomy followed by ultrasound stimulation and imaging under anesthesia [7], [10], [11]. However, this design can be confounded by factors such as surgical stress from the craniotomy, variability in procedure time (and thus anesthetic duration), and inflammation in the mouse brain [16], [17]. Additionally, isoflurane and other commonly used anesthetic agents lead to vasodilation and disruptions in neurovascular coupling [18], which can interfere with ultrasound stimulation effects [19]. Recognizing these issues, Lecea’s group developed the Fiber Photometry Coupled Focused Ultrasound System (PhoCUS) to investigate ultrasound stimulation in freely behaving mice [20], [21]. However, this approach requires inserting an optical fiber probe in mice brain, which increases invasiveness and may potentially interfere with ultrasound stimulation. Other groups have used a craniotomy-based glass cranial window to image the mouse brain in a head-fixed, awake state [19], [22]. However, craniotomy procedures can lead to microglia activation and inflammation [17]. Moreover, the use of a glass window in awake studies can interfere with ultrasound stimulation due to acoustic impedance mismatches, thereby limiting the potential for coaxial ultrasound stimulation and optical recording. Another approach utilized a clear intact skull preparation [14], [23]. While this approach reduces inflammatory confounds, the intact skull window is limited in resolution and may not support high-resolution imaging techniques, such as two-photon microscopy, due to skull aberrations.

A straightforward solution to resolve the shortcomings is to use transparent ultrasound transducers (TUTs), which allows coaxial optical illumination and ultrasonic transduction and reception. Since our first demonstration of lithium niobate-based piezoelectric TUTs for photoacoustic imaging in year 2019 [24], there has been considerable interest in exploring different biomedical application of LN-TUTs [25], [26]. In particular, our group has first demonstrated that a miniaturized lithium niobate-based TUT can be implanted as a thinned-skull cranial window (wearable ultrasound transducer) to allow multimodal photoacoustic and optical neuroimaging in awake mice [27]. Beyond photoacoustic imaging, our group also previously demonstrated TUT as an on-chip platform for stimulating cancer cells directly cultured on the biocompatible TUT surface [28]. However, TUTs remain largely unexplored for investigating ultrasound neuromodulation. Lee et al. [29] demonstrated an acousto-optic window using a PVDF-TrFE based ultrasound transducer. However, the window had low optical transparency (∼40%) and acoustic pressure output (∼110 kPa). Additionally, their study involved craniotomy, and the mice were imaged and stimulated in anesthetized states, thereby suffering from the same confounds mentioned previously.

Building on the strengths of our prior work that demonstrated awake mouse brain imaging using TUTs [27], in this study we investigate the potential of TUT cranial window for ultrasound neuromodulation and simultaneous optical imaging. We use laser speckle contrast imaging (LSCI) and intrinsic optical signal imaging (IOSI) to investigate cortical hemodynamic changes induced by ultrasound stimulation through TUT cranial window in awake mice. Furthermore, by integrating the TUT with a thinned-skull preparation and head restraint awake mice imaging, we mitigate confounding factors such as anesthesia and inflammation. Our results demonstrate that the TUT cranial window enables the generation of ultrasound stimuli with sufficient acoustic pressure capable of eliciting cortical hemodynamic changes and facilitates imaging of multiple contrasts (speckle contrast and optical absorption) across different optical wavelengths. Furthermore, we observed that ultrasound stimulation induces increases in both cerebral blood flow (CBF) and total hemoglobin (HbT). By systematically varying ultrasound parameters, we showed that peak and cumulative hemodynamic changes are directionally correlated with the total stimulation time and ultrasonic intensity. Overall, these findings underscore the potential of TUTs as valuable tools for researchers investigating the mechanisms underlying ultrasound neuromodulation.

## 2 Methods

### 2.1 TUT fabrication and characterization

The TUT fabrication followed the process mentioned previously in [27], [30], see supplemental methods for fabrication details. A 4×4 mm^2^ TUT was fabricated with 3×3 mm^2^ lithium niobate element. The TUT then underwent electrical, pressure and thermal characterization, see supplemental methods for details.

### 2.2 TUT cranial window preparation

For implantation, we followed a procedure similar to that outlined by [16], [27]. In summary, three C57BL/6J mice (male, 2–6 months old, Jackson Laboratory, Bar Harbor, ME, USA) were used for the thinned-skull TUT cranial window preparation. The surgical procedure involved fixing the mouse on a stereotaxic apparatus and creating a midline incision in the skin to expose the skull. A high-speed drill was used to thin the skull over an approximately 4 × 4 mm^2^ region around the somatosensory cortex until the area was transparent. Precautions were taken to periodically flush the drilled skull with cold saline to reduce heating. The TUT was implanted over the thinned region using a thin layer of cyanoacrylate glue. A headbar was attached to the base of the skull with cyanoacrylate glue and dental cement. The remaining exposed skull was then also covered with glue and dental cement. The implanted TUT cranial window is shown in Fig. 1. The mice were subsequently monitored for one week to recover from surgery and then incrementally acclimated to the head bar mount for three days. The image of TUT cranial window and awake head restrained mice on head bar mount is shown in Fig. 1. For anesthesia recording, the mice were administered isoflurane mixed with medical oxygen at 1L/min using a nose cone. All animal procedures were carried out in accordance with the Institutional Animal Care and Use Committee (IACUC) guidelines at Pennsylvania State University.

**Figure 1:**
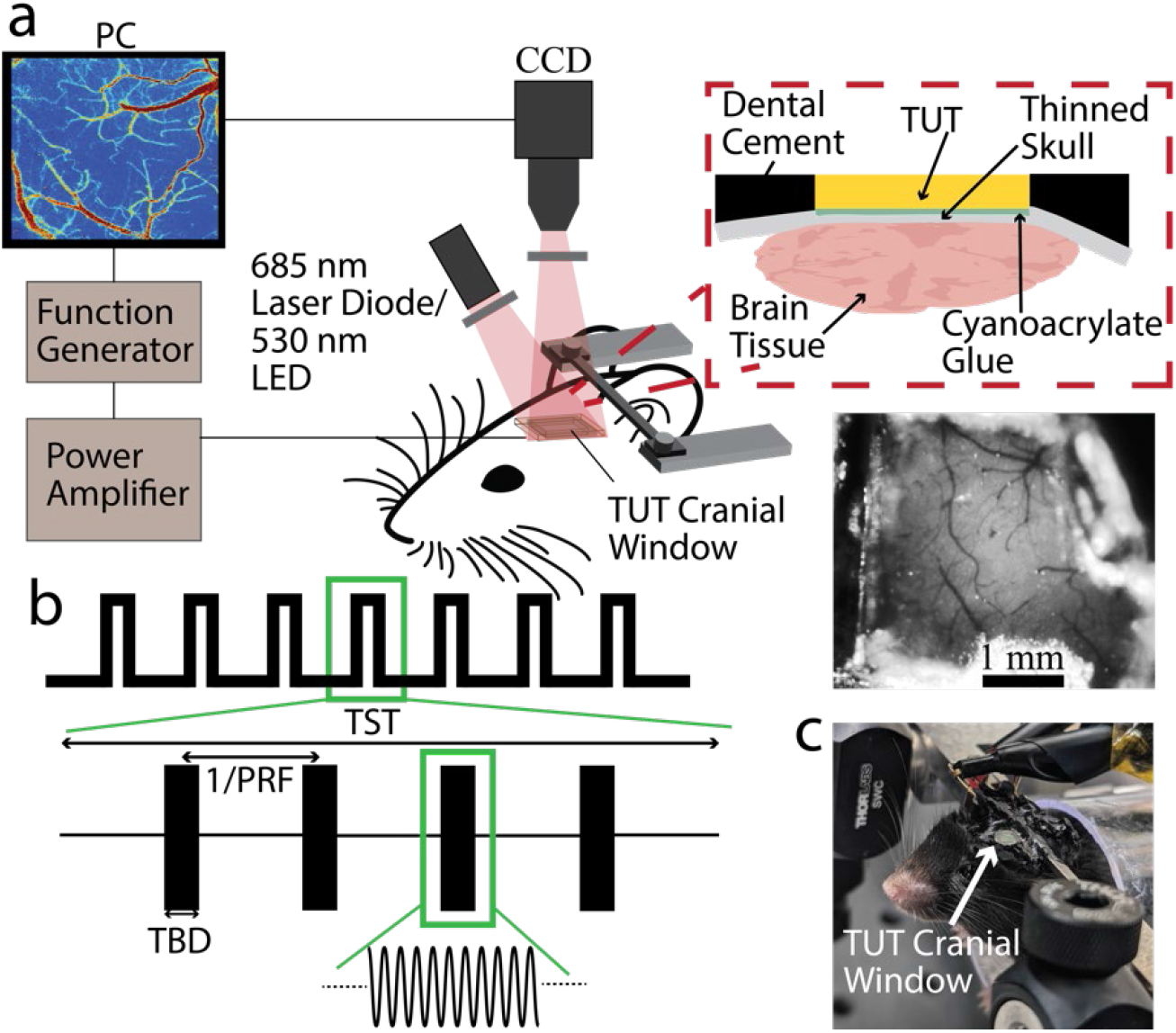
TUT based optical imaging ultrasound stimulation setup: (a) Schematic of the optical imaging setup combined with ultrasound stimulation on awake mouse brain; red inset represents close-up cross-sectional view of TUT thinned skull cranial window; and image below displays an actual image of the TUT cranial window, with vessels visible through the thinned skull and TUT. (b) the stimulation paradigm used in our setup, each experiment consisted of seven trials, thirty seconds apart. (c) the photograph of mice on the head mounted setup for optical imaging in awake state. TUT: Transparent ultrasound transducer, TST: Total stimulation time, PRF: Pulse repetition frequency, TBD: Total burst duration.

### 2.3 Optical imaging and ultrasound neuromodulation

The optical imaging setup is illustrated in Fig. 1a and was based on [31], [32], [33]. A 685 nm laser diode (HL6750MG, Thorlabs, NJ, USA) was used for LSCI, and a 530 nm light-emitting diode (M530L4, Thorlabs, NJ, USA) was used for IOSI. The laser light passed through a plano-convex lens to ensure uniform illumination of the brain tissue through the transparent ultrasound transducer. Two polarizers were arranged in a cross-polarization configuration in illumination and detection paths to minimize specular reflections for LSCI. The light was collected using a Nikon AF Micro Nikkor 60 mm f/2.8D lens (Nikon, Tokyo, Japan) and recorded using a CCD camera (Infinity 1RM, Teledyne Lumenera, ON, Canada) with a frame rate of 27.4 frames per second and exposure times of 20 ms for LSCI and 100 ms for IOSI. Each recording consisted of seven ultrasound stimuli spaced thirty seconds apart, with a total recording time of 240 seconds per ultrasound parameter, capturing a 600×480 pixel area corresponding to an approximately 3 × 3 mm^2^ region. A custom MATLAB script was used to control the camera acquisition and trigger ultrasound pulses at the desired times. The imaging was conducted consecutively. For ultrasound stimulation, the TUT was excited using an electrical pulse train produced by a function generator (SDG6022X, Siglent, Ohio, USA) and amplified with a 50 dB RF amplifier (310L, E&I, NY, USA). Table S1 summarizes the various ultrasound parameters used in different sets of experiments, and the pulse pattern is depicted in Fig. 1b.

### 2.4 Data analysis

LSCI infers blood flow by analyzing variations in speckle patterns caused by the movement of red blood cells. Speckle contrast, defined as the ratio of the standard deviation (*σ*) of pixel intensity to the mean intensity (μ), over either a spatial or temporal window, is given by:

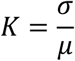

For Lorentzian flow, speckle contrast is related to the decorrelation time *τ*_*c*_ and the camera exposure time *T* through the formula:

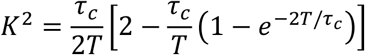

When the decorrelation time is much shorter than the exposure time (typical in blood flow measurements), the relationship simplifies to:

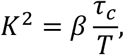

where *β* is a factor accounting for system coherence and averaging effects. Since blood flow is inversely proportional to the decorrelation time, it can be inferred as being inversely proportional to 1*/K*^2^ and was used therefore to obtain CBF maps in our study. For our analysis, we used spatial LSCI with spatial window of 5. The LSCI image analysis was done in MATLAB with scripts mentioned in [34], [35] and were further modified for our study. We further generated ROI based pixel-by-pixel relative CBF maps based on the following formula for relative cerebral blood flow (% ΔCBF) changes:

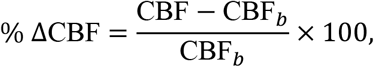

where CBF_*b*_ represents the baseline CBF.

The IOSI signal is generated by optical absorption and subsequent changes in the optical density of light diffusing back to the camera. The optical density changes (ΔOD) were computed as:

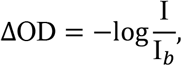

where *I* is the light intensity and *I*_*b*_ is the baseline light intensity. Furthermore, as we used 530 nm wavelength LED for IOSI which is in close proximity to oxy- and de-oxyhemoglobin isosbestic point, the ΔOD changes were inferred as HbT changes. To process the recorded videos, we matched the ROI selection to the ROI used in LSCI processing and subsequently generated pixel-by-pixel ΔOD maps [36]. For both LSCI and IOSI, a two-second baseline was considered before each trial, and all trial curves were averaged together. A moving average window of (% ΔCBF) was used for LSCI traces to reduce high frequency fluctuations.

## 3 Results

### 3.1 TUT characterization results

We first present the characterization results of the TUT. A picture of the TUT is shown in Fig. 2a indicating its transparent nature. As demonstrated in our previous studies, TUTs exhibit significant optical transparency and are viable for optical imaging [27]. Next, we characterized the impedance and phase curves of the TUT in Fig. 2b. The 36^°^ Y-cut lithium niobate element has a thickness of 250 μm and a sound speed of 7238 m/s along the Z-axis [37]. Given the resonance frequency (*f*_*r*_) is calculated as:

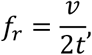

where *v* is the speed of sound and *t* is the thickness of the lithium niobate crystal, we expect a resonant frequency of 14.4 MHz. However, with the addition of the Parylene-C acoustic matching layer, two resonant peaks (11.6 and 15.4 MHz) and two anti-resonant peaks (11.9 and 15.7 MHz) are observed. To identify the frequency for maximum pressure output, we characterized the pressure output of the TUT submerged in deionized water at different frequencies using a bullet hydrophone. The results indicate that the TUT achieves maximum response in the 12–13 MHz frequency band (Fig. S1a). Therefore, we fixed the ultrasonic stimulation frequency at 12 MHz for this study. Fig. S1a further shows that the TUT can be driven at different frequencies, albeit at reduced pressures.

**Figure 2:**
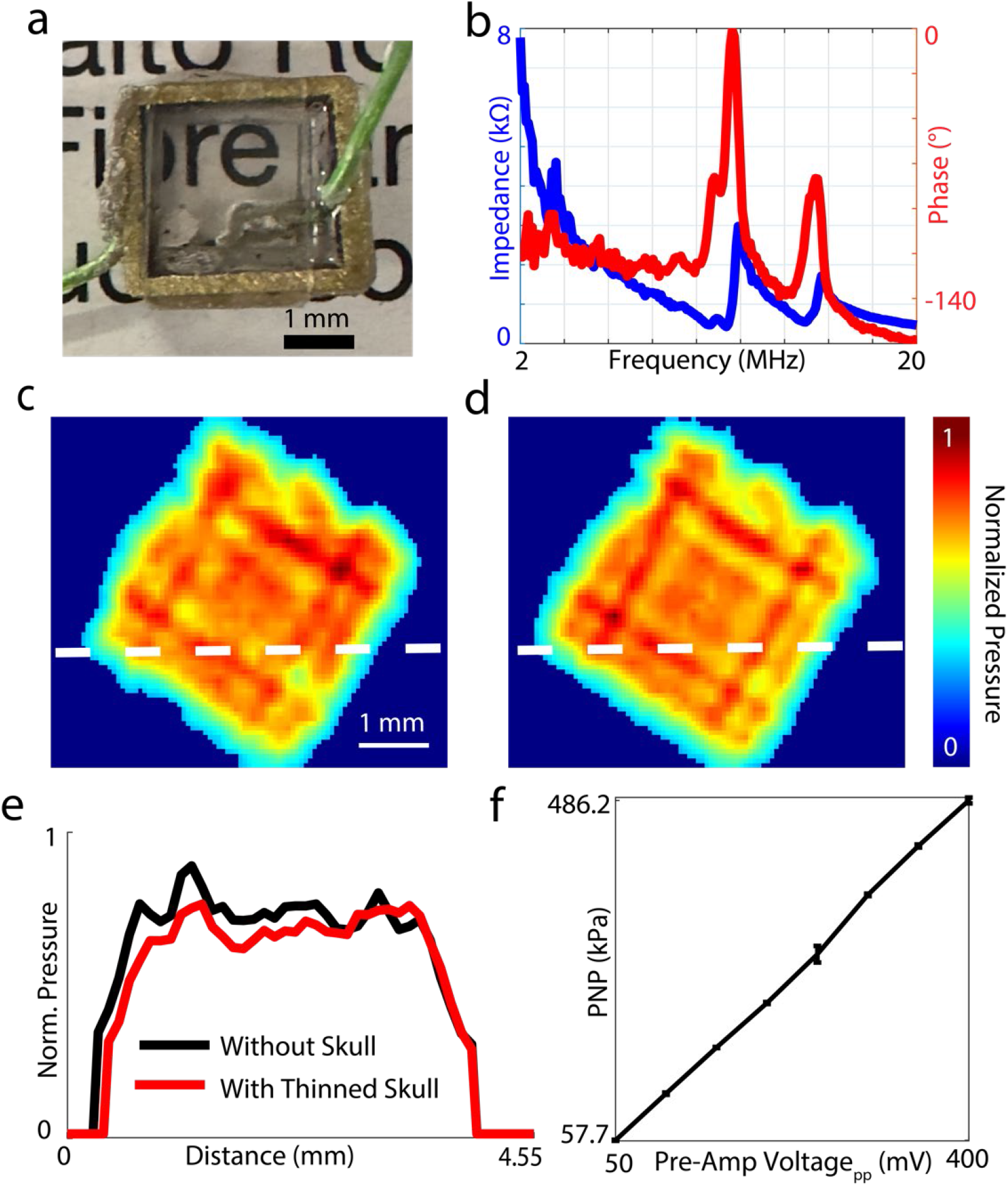
TUT characterization: (a) a picture of the fabricated TUT, highlighting its transparent nature; (b) displays the impedance and phase curves of the TUT; (c, d) the normalized pressure output with and without a thinned skull; (e) presents the spatial plot of normalized pressure output along the dashed line for both cases in (c,d); (f) depicts the peak negative pressure (PNP)output of the TUT at various voltage inputs.

Next, we characterized the acoustic pressure response across the TUT surface (Fig. 2c) at 1.5 mm distance. The pressure distribution is uniform, and we also demonstrated that the thinned skull causes minimal distortion to the ultrasound signal (Fig. 2d, e). However, given that our TUT cranial window preparation involved the use of cyanoacrylate glue for the reinforcement of the thinned skull, we observe a 30% peak pressure drop with its addition. Consequently, we derated our pressure values by 30% to account for the same (Table S1). We also examined the pressure attenuation in mouse brain tissue at 12 MHz (Fig. S1b). We observed a 6 dB pressure attenuation for 4.2 mm of mouse brain tissue. These results align with tissue attenuation values reported in the literature at 9.84 dB/cm for 12 MHz [38]. Lastly, we characterized the derated peak negative pressure (PNP) output of the TUT at different input voltages, showing that the TUT exhibits a linear response to increasing voltages (Fig. 2f). Subsequently, we calculated the *I*_SPPA_ and MI for the various stimulation parameters listed in Table S1. The maximum *I*_SPPA_ was 4.9 W/cm ^2^, and the maximum MI was 0.12. Additionally, we looked at two potential sources of thermal effects: ultrasonic absorption and heat dissipation from TUT surface. We performed numerical calculations of the maximum temperature rise in tissue due to ultrasonic absorption. The maximum temperature rise in tissue was found to be 0.03 ^°^C at PNP: 486.2 kPa, PRF: 1 kHz, TBD: 500 *μ*s, and TST: 250 ms. Under the same ultrasound parameters, the maximum temperature rise across the surface of the TUT was found to be 1.3 ^°^C (Fig. S2). Overall, the temperature rise was transient, given the maximum stimulation duration of 250 ms, and remains below the temperature damage threshold for the mouse brain [39].

### 3.2 TUT cranial window is feasible for ultrasound neuromodulation

Next, we investigated the feasibility of our TUT cranial window in eliciting hemodynamic responses in response to ultrasound stimulation using two modalities: LSCI for measuring % ΔCBF and IOSI to estimate Δ OD changes. The choice of these two modalities was motivated by three key reasons: (i) to understand the interplay between CBF and HbT changes, (ii) to demonstrate that the TUT cranial window can be used to image the rodent brain at different wavelengths, and (iii) to leverage distinct imaging contrasts (speckle changes and optical absorption) to obtain cortical hemodynamic information. Fig. 3a, b, and supplementary videos 1, 2 show that ultrasound stimulation (PRF: 1 kHz, TBD: 500 *μ*s, PNP: 486.2 kPa, TST: 250 ms) induced increases in both CBF and HbT. Fig. 3c displays the averaged % ΔCBF and ΔOD traces across mice, with their peaks occurring at 1.5 s and 1.7 s, respectively, and both returning to baseline by 4 s post ultrasound stimulation. However, in the % ΔCBF trace, an additional peak is observed at 0.9 s. We hypothesized that this might result from increased static speckle interference from within the TUT and surrounding tissue following ultrasound stimulation. To confirm this, we fixed the TUT in a 1% v/v intralipid phantom to evaluate its impact when aligned with the optical imaging paths (Fig. S3a). As expected, in LSCI, the intralipid phantom also exhibited a % Δflow peak at 0.9 s when subjected to the same ultrasound stimulus (PRF: 1 kHz, TBD: 500 *μ*s, PNP: 486.2 kPa, TST: 250 ms, Fig. S3b). The slower decay to baseline in the phantom compared to the in vivo study may be attributed to differences in the elasticity of the intralipid medium causing the delay. Conversely, IOSI of the intralipid phantom showed minimal changes in ΔOD under the same ultrasound stimulus Fig. S3c. This finding indicates two key points: (i) ultrasonic vibrations caused static speckle changes, leading to the initial rise in % ΔCBF, and (ii) IOSI measurements experience minimal interference from ultrasonic vibrations. Consequently, to validate our CBF signal, we performed a one-second moving window correlation between the averaged CBF and IOSI traces. Soon after the ultrasound stimulus, the correlation coefficient dropped to 0.1 but eventually recovered post stimulation after 1 s post stimulation Fig. 3d. Therefore, to ensure minimization of potential biases from speckle interference, all analysis was done starting 1.2 s after the ultrasound stimulus.

**Figure 3:**
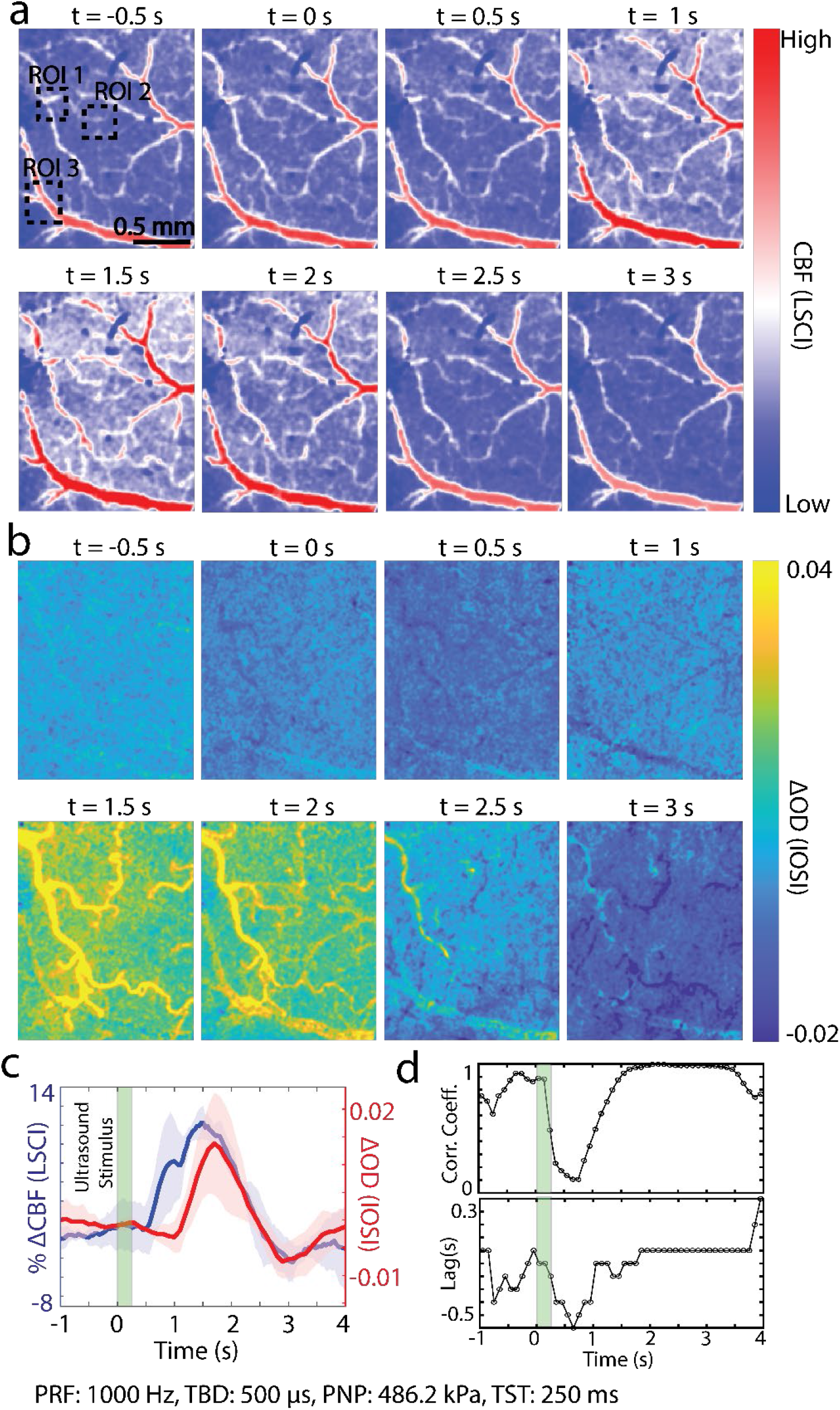
Ultrasound stimulation induced CBF and HbT (ΔOD) changes in the brain, (a) the LSCI CBF maps and (b) the IOSI ΔOD maps with ultrasound stimulation at t = 0; (c) the overlaid averaged % ΔCBF and ΔOD trials (n = 7) with shaded region indicating the SEM for a single mice; (d) the correlation coefficient and time lag calculated over a one second moving window between % ΔCBF and ΔOD traces.

### 3.3 Multiparametric investigation of ultrasound neuromodulation

Ultrasound stimulation involves a multi-dimensional parameter space, requiring careful investigation of different parameter sets. The parameter set indicated in Table S1 was used to examine the relationship between ultrasound-induced hemodynamic changes and ultrasound stimulation parameters including TST, PNP, total burst duration (TBD) and pulse repetition frequency (PRF). To perform a statistical investigation, three mice underwent LSCI and IOSI imaging sessions. The trial-averaged traces for each mouse from each session were combined and are presented in Fig. 4 and Fig. 5. Fig. 4a1, b1 show the LSCI and IOSI traces, and their corresponding mean peak and cumulative (area under the curve, AUC) changes are plotted in Fig. 4a2-3, b2-3. These data illustrated the effects of increasing stimulation time from 25 to 250 ms at PRF: 1000 Hz, TBD: 500 *μ*s, and PNP: 486.2 kPa. We observed that increasing the stimulation time results in a significant increase in both the peak change and AUC for % ΔCBF and ΔOD, with stimulation times of 200 ms and 250 ms showing significant differences compared to 25 ms and 50 ms (Fig. 4a2-3, b2-3, Mean±SEM, Friedman test, *p<0.05, **p<0.01). Similarly, in Fig. 4c, d, we examined the effect of increasing PNP from 230.8 to 486.2 kPa at PRF: 1000 Hz, TBD: 500 *μ*s, and TST: 250 ms. This resulted in a monotonic increase in % ΔCBF and ΔOD. Significant differences were observed between PNP values of 486.2 kPa and 230.8 kPa for both the peak change and AUC of % ΔCBF and ΔOD (Fig. 4c2-3, d2-3, Mean±SEM, Friedman test, *p<0.05). These results follow expected trends, with larger ultrasound doses eliciting greater hemodynamic changes. Additionally, in Fig. 5, we investigate the effects of changing the TBD and PRF. When varying TBD from 100 to 500 *μ*s at PRF: 1000 Hz, TST: 250 ms, and PNP: 486.2 kPa, we only observe significant changes in the peak and AUC of % ΔCBF when increasing TBD from 100 to 500 *μ*s. (Fig. 5a2-3, Mean±SEM, Friedman test, *p<0.05). For ΔOD, both the peak and AUC changes showed significant differences between 200 and 400 *μ*s while minimal differences are observed between other parameters (Fig. 5b2-3, Mean±SEM, Friedman test, *p<0.05). Lastly, we examined changes in PRF from 100 to 1000 Hz at TBD: 500 *μ*s, TST: 250 ms, and PNP: 496.1 kPa. No clear trends are observed in the peak or AUC of % ΔCBF. Conversely, ΔOD showed significant increases in both the peak and AUC between 100 Hz and 300 or 500 Hz (Fig. 5c2-3, b2-3, Mean±SEM, Friedman test, *p<0.05).

**Figure 4:**
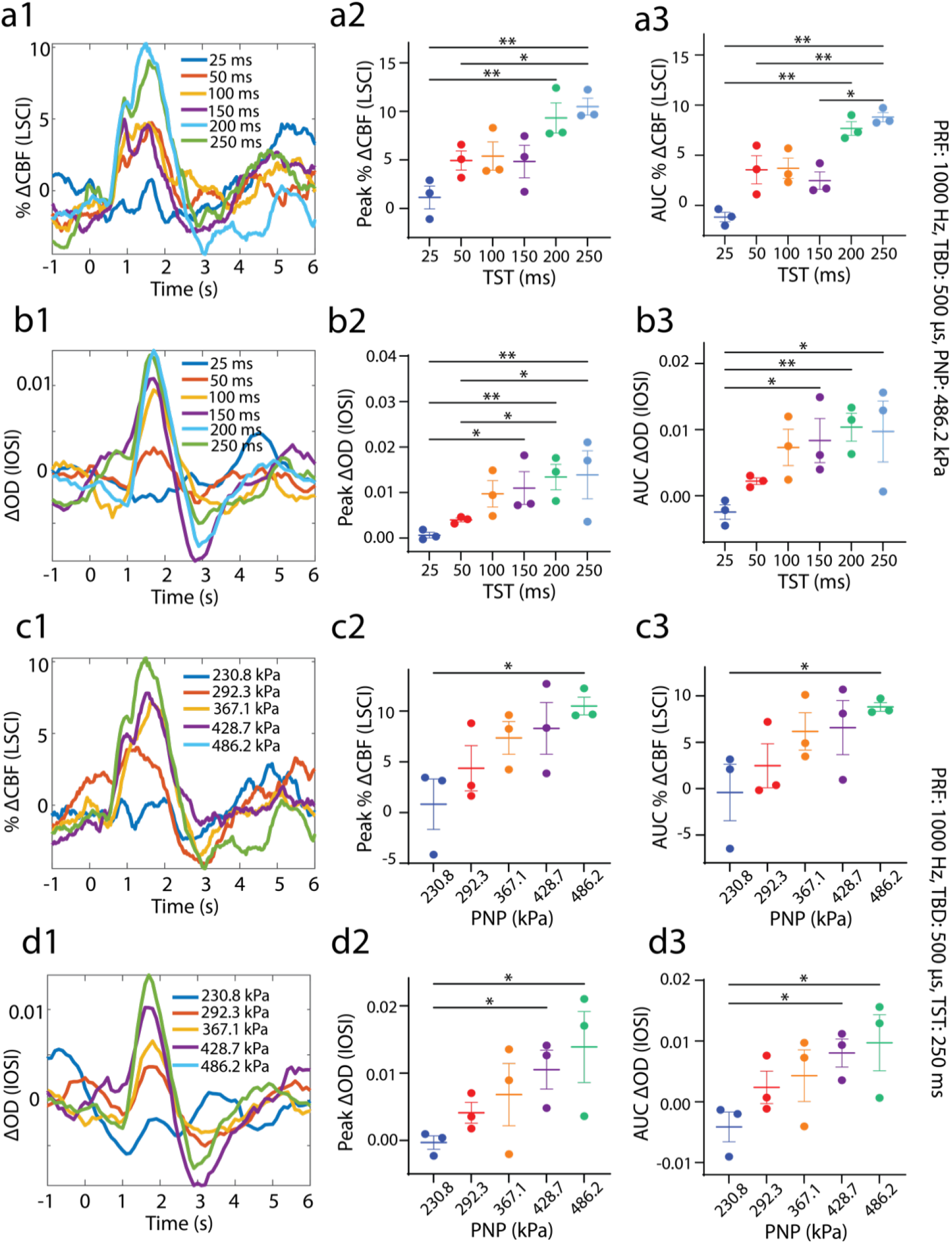
Relationship between ultrasound TST and PNP and elicited hemodynamic responses: Averaged % ΔCBF traces across different mice in response at different ultrasound TST (a1) and PNP (c1). Statistical comparison for peak and area under curve (AUC) % ΔCBF change at different ultrasound TST (a2, a3) and PNP (c2, c3). Averaged ΔOD traces across different mice in response at different ultrasound TST (b1) and PNP (d1). Statistical comparison for peak and AUC ΔOD changes at different ultrasound TST (b2, b3) and PNP (d2, d3). (n=3, Mean±SEM, Friedman test, *p<0.05, **p<0.01) TST: Total stimulation time, PRF: Pulse repetition frequency, TBD: Total burst duration.

**Figure 5:**
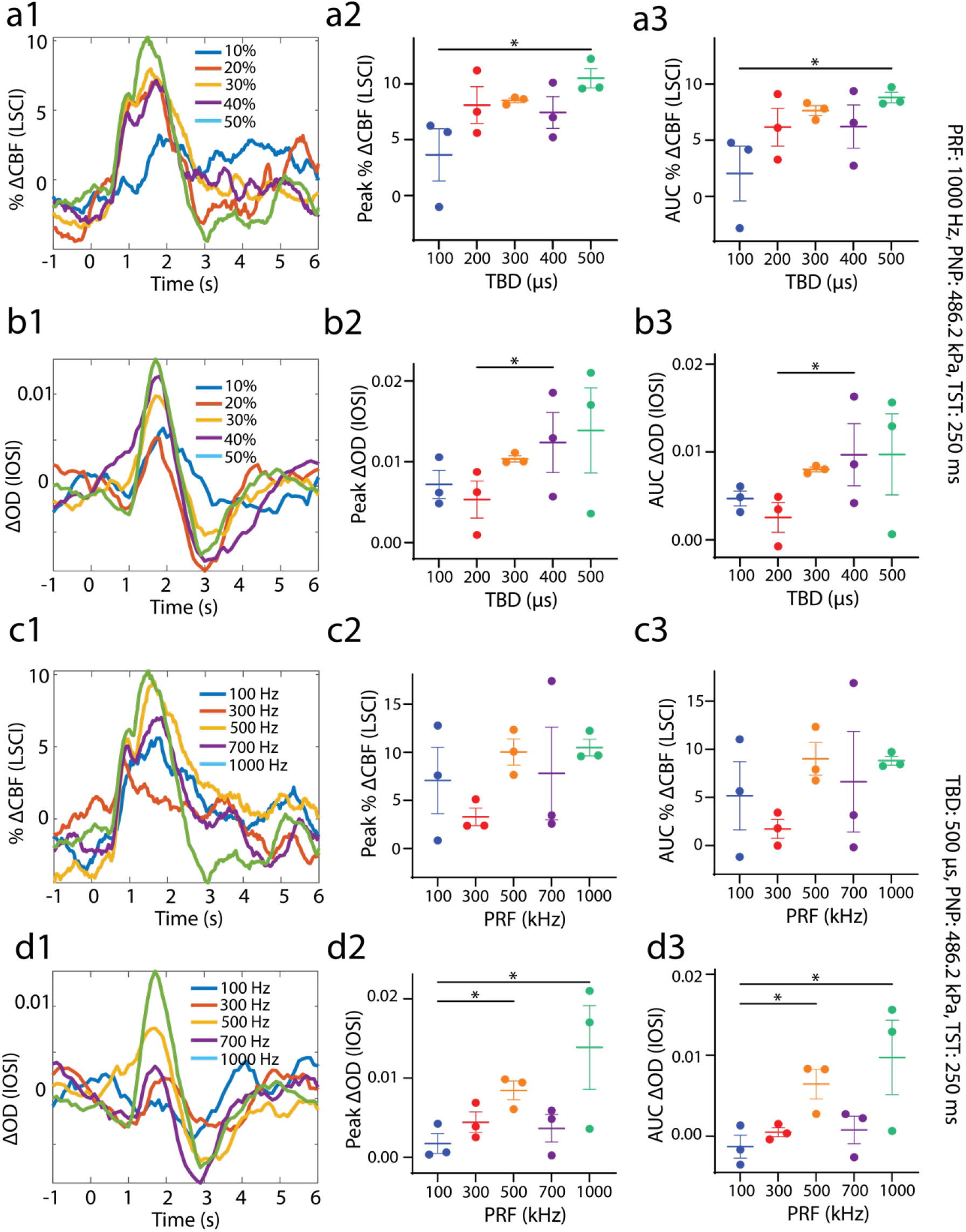
Relationship between ultrasound TBD and PRF and elicited hemodynamic responses: Averaged % ΔCBF traces across different mice in response at different ultrasound TBD (a1) and PRF (c1). Statistical comparison for peak and area under curve (AUC) % ΔCBF change at different ultrasound TBD (a2, a3) and PRF (c2, c3). Averaged ΔOD traces across different mice in response at different ultrasound TBD (b1) and PRF (d1). Statistical comparison for peak and AUC ΔOD change at different ultrasound TBD (b2, b3) and PRF (d2, d3). (n=3, Mean±SEM, Friedman test, *p<0.05) TST: Total stimulation time, PRF: Pulse repetition frequency, TBD: Total burst duration.

## 4 Discussion

In this study, we proposed a TUT-based thinned-skull cranial window to investigate ultrasound neuromodulation in awake mice. Our findings demonstrated that the in-house-fabricated TUT produces ultrasonic outputs sufficient for eliciting ultrasonic neuromodulation. By employing the TUT as a thinned-skull cranial window, we seamlessly integrated pathways for both optical and ultrasound stimulation, enabling investigations of ultrasound neuromodulation in the awake mouse brain. Using LSCI, we showed that our stimulation paradigm induced an increase in ΔCBF following ultrasound stimulation. These results are further substantiated by IOSI, where we observed an increase in ΔOD indicative of elevated HbT. Furthermore, through a multiparametric analysis, we demonstrated that ΔCBF and ΔOD are directionally correlated to TST and PNP. However, while increased ΔCBF and ΔOD are observed, no dependent trend is noted w.r.t changing TBD or PRF.

There have been few studies detailing the hemodynamic changes in response to ultrasound stimulation in vivo; a comparison of different studies in literature with ours is presented in Table S2. Our results are consistent with existing studies, despite the use of a higher frequency (12 MHz) ultraosund transducer. Yuan et al. [7] used a 0.5 MHz transducer to show that ΔCBF increased in anesthetized mice 2.3 seconds after ultrasound stimulation. The monotonic elevation in ΔCBF with TST and PNP, along with the lack of observable differences with changes in duty cycle (or TBD), aligns with our findings. Another study using IOSI in awake mice [19] observed a rapid increase in cortical hemodynamics within a similar timescale (<2 seconds) as reported here. Furthermore, they also showed that peak hemodynamic responses were reduced and delayed with increases in anesthetic levels. We further validated the same with our setup (Fig. S4) where we see hemodynamic changes are attuned and exhibit delayed onset with increased anesthesia. Thereby, showing the importance of awake mice imaging to overcome anesthetic biases that affect ultrasound neuromodulation. More recent studies from Konofagou’s group [10], [11] employed functional ultrasound to investigate cerebral blood volume changes in anesthetized mice and non-human primates. Using stimulation durations of several seconds and a 4 MHz transducer probe in vivo, they also reported an increase in cerebral blood volume in mice. Interestingly, in our study, changing PRF did not result in an observable trend. This may be attributed to previously reported intrinsic cell specificity to ultrasound stimuli, where inhibitory and excitatory neurons respond differently to PRF, as demonstrated by [6], [40]. Regional differences in tissue elasticity between cortical and subcortical areas [11], [41] could also contribute to these observations. A recent study by Murphy et al. [20] employed a 0.55 MHz ring ultrasound transducer to identify stimulation parameters, showing that low PRF can lead to both inhibition and excitation. These studies are particularly relevant to our work, as the hemodynamic changes observed at different PRF may reflect an interplay between inhibitory and excitatory mechanisms, warranting further investigation.

Our approach of using a TUT cranial window offers several distinct advantages. A key aspect of our study is the use of a custom-made TUT, which can be scaled to different sizes and frequencies based on the application. Although the TUT is unfocused, its dimensions (3×3 mm^2^) allow it to produce ultrasound stimulus spot sizes comparable (and even smaller) to those generated by focused ultrasound transducers operating at frequencies below 1 MHz (Table S2). Another significant advantage of the TUT thinned-skull cranial window is that it is integrable in conventional glass based cranial window recording setups. Furthermore, our reinforced thinned-skull preparation reduced glial activation otherwise associated with the craniotomy procedure [16]. Additionally, the TUT cranial window on account of being reinforced with cyanoacrylate glue is stable for more than 10 weeks (Fig. S5) and can therefore potentially enable longitudinal imaging of ultrasound neuromodulation effects in healthy and diseased mice models. Lastly, TUTs fabrication is inexpensive (<20$) in comparison to conventionally used ultrasound transducers with costs >1000$. Moreover, the fabrication process is easily accomplished with standard fabrication tools commonly available to researchers across universities. Thereby making TUTs an inexpensive and accessible alternative to conventional ultrasound transducers.

## 5 Conclusion

In conclusion, we present a novel TUT cranial window platform for investigating ultrasound neuromodulation in awake mice. This platform enables coaxial imaging and ultrasound stimulation, with cortical hemodynamic responses that align with existing literature, validating the feasibility of the TUT cranial window. The proposed technique significantly simplifies ultrasound stimulation setups and can be easily integrated into laboratories already using glass-based cranial windows. Additionally, the custom-made TUTs are flexible, allowing scalability in size and frequency. In the future, by utilizing thicker lithium niobate crystals, we aim to develop a lower-frequency TUT cranial window capable of accessing deeper brain regions. Furthermore, by interleaving various wavelengths, we plan to perform simultaneous multimodal optical imaging in the future. Lastly, the TUT can also be scaled to a transparent transducer array that will allow for steering ultrasound neuromodulation to different regions in the brain while enabling avenues for multimodal ultrafast ultrasound, photoacoustic and optical imaging [42], [43]. Overall, the proposed TUT cranial window overcomes technical challenges in integrating ultrasound stimulation with optical imaging and offers a robust and a promising tool to elicit ultrasound neuromodulation, thereby opening avenues for mechanistic and therapeutic investigations of ultrasound neuromodulation in healthy and diseased rodent models.

## Author contributions

**Shubham Mirg**: Conceptualization, Methodology, Software, Data curation, Visualization, and Investigation; **Krishnendu Samanta**: Investigation and Methodology; **Haoyang Chen**: Methodology; **Jin Jiang**: Data curation; **Kevin L. Turner**: Methodology; **Fatmeh Salehi**: Methodology; **Kathiravan M. Ramiah:** Data Curation**; Patrick J. Drew**: Methodology, Resources, and Investigation; **Sri-Rajasekhar Kothapalli**: Conceptualization, Data curation, Methodology, Supervision, and Investigation. Shubham Mirg wrote the original draft and co-authors provided comments and edited the manuscript.

## Data availability

The raw data supporting the findings of this study are available from the corresponding author upon reasonable request. The processed data used in the analysis have been deposited in zenodo and can be accessed at https://doi.org/10.5281/zenodo.14876675.The code for generating the figures is available on GitHub at https://github.com/shubhammirg/TUTCranialWindowStim.

