## Supplementary File for "Integrated Ultrasound Neuromodulation and Optical Neuroimaging in Awake Mice using a Transparent Ultrasound Transducer Cranial Window"

### Methods

### TUT fabrication

For TUT fabrication, we follow the process described in [1], [2]. In brief, 200 nm of indium tin oxide (ITO) was coated onto 250 $\mu$m lithium niobate crystals (36° Y-cut, Precision Micro Optics, Burlington, MA, USA). Subsequently, the ITO-coated lithium niobate crystal was diced into 3 mm $\times$ 3 mm pieces. A coaxial wire was connected using silver epoxy (8330S, MG Chemicals, Ontario, Canada), and a brass housing was placed around the crystal. Transparent epoxy (EPO-TEK 301, Epoxy Technology, MA USA) was poured over the lithium niobate crystal within the brass housing to act as a backing layer. A cover glass slip was placed on top to provide stability. After the epoxy was cured, another coaxial wire was connected to the brass tube using silver epoxy, and the brass tube was then connected to the ground layer of the crystal, also using silver epoxy. Lastly, a 30 $\mu$m coat of Parylene-C was added for acoustic impedance matching, ensuring biocompatibility, and providing electrical isolation. The overall thickness of TUT was 1.2 mm.

### TUT characterization

A network analyzer (E5100A, Agilent, CA, USA) was used to analyze the impedance of the TUT to obtain the resonant and anti-resonant peaks. The pressure field measurement and characterization were conducted using a pre-calibrated capsule hydrophone (AH-2010-100, ONDA, California, USA). The pressure output was recorded by positioning the TUT coaxially with the hydrophone and exciting the TUT at different voltage and frequency inputs. The effect of cyanoacrylate glue on pressure output was also evaluated by applying a thin layer of glue over the TUT. Freshly harvested brain tissue of varying thickness was placed on the transducer with the help of 1.5% agarose, and the peak pressure output was recorded to estimate tissue attenuation. For pressure field mapping, the hydrophone was scanned laterally across the surface of the transducer using a linear stage at 1.5 mm distance from the TUT. To understand the effects of a thinned mouse skull, a $3\times3 \mathrm{mm}^{2}$ piece of thinned skull was placed on the TUT and secured with 1.5% agarose, and the hydrophone scanning was repeated. Following this, the spatial peak pulse average intensity ($I_{\text{SPPA}}$) for different parameters was calculated as:

$$I_{\text{SPPA}}=\frac{1}{\text{PD}}\int\frac{p^{2}\left( t \right)}{Z_{0}} dt,$$

where $\text{PD}$ is the pulse duration, $p\left( t \right)$ is the instantaneous temporal pressure profile at the spatial maximum, and $Z_{0}$ is the characteristic acoustic impedance given by $\rho c$, where $\rho$ is the density of the medium, and $c$ is the speed of sound in the medium. We used $\rho=1040 kg/m^{3}$ and $c=1560 m/s$ for mouse brain tissue. Furthermore, the mechanical index (MI) was calculated using:

$$MI=\frac{\text{PNP}}{\sqrt{f}},$$

where $\text{PNP}$ is the peak negative pressure, and $f$ is the ultrasound frequency. We identified two potential heat sources: one generated through ultrasonic absorption in the tissue and another generated at the surface of the TUT. To evaluate the thermal effects of ultrasonic absorption, the maximum temperature rise in the tissue was numerically calculated using [3]:

$$\Delta T=\frac{Q\cdot\text{TST}}{C_{v}},$$

where

$$Q=2\alpha\times I_{\text{SPTA}},$$

and $\text{TST}$ is the total stimulation time (the total exposure time of the tissue to ultrasound), $C_{v}$ is the heat capacity per unit volume ($3.83 J/cm^{3}/^{\circ}C$ for brain tissue [4]), $\alpha$ is the attenuation in tissue ($9.84 dB/cm$ at 12 MHz [5]), and $I_{\text{SPTA}}=I_{\text{SPPA}}\times\text{Duty Cycle}$. To measure the temperature, rise at the surface of the TUT, we used a K-type point thermometer (T3000FC, Fluke Corporation, WA, USA) placed on the surface of the TUT with a layer of cyanoacrylate glue, surrounded by an acoustic gel coupling medium.

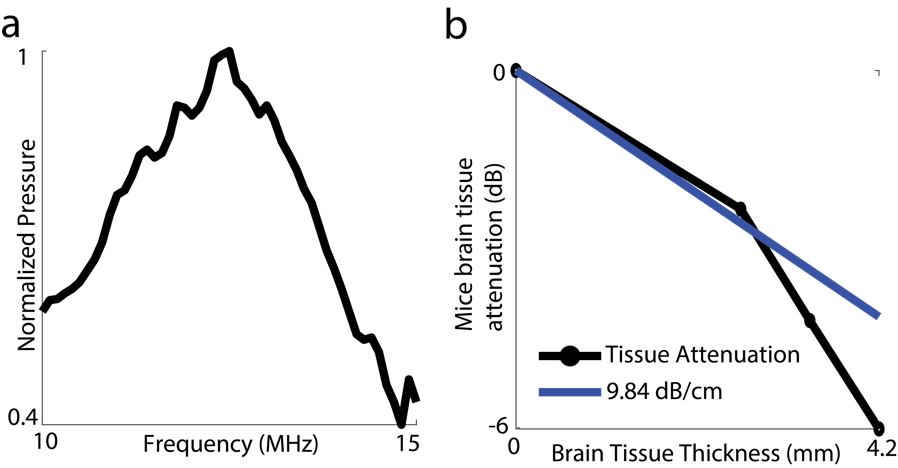

Figure S1: a) the normalized pressure output at different frequencies, with 12-13 MHz frequency band exhibiting >90% of peak pressure. b) the tissue attenuation across mice brain in comparison to the expected tissue attenuation of 9.84 dB/cm [5].

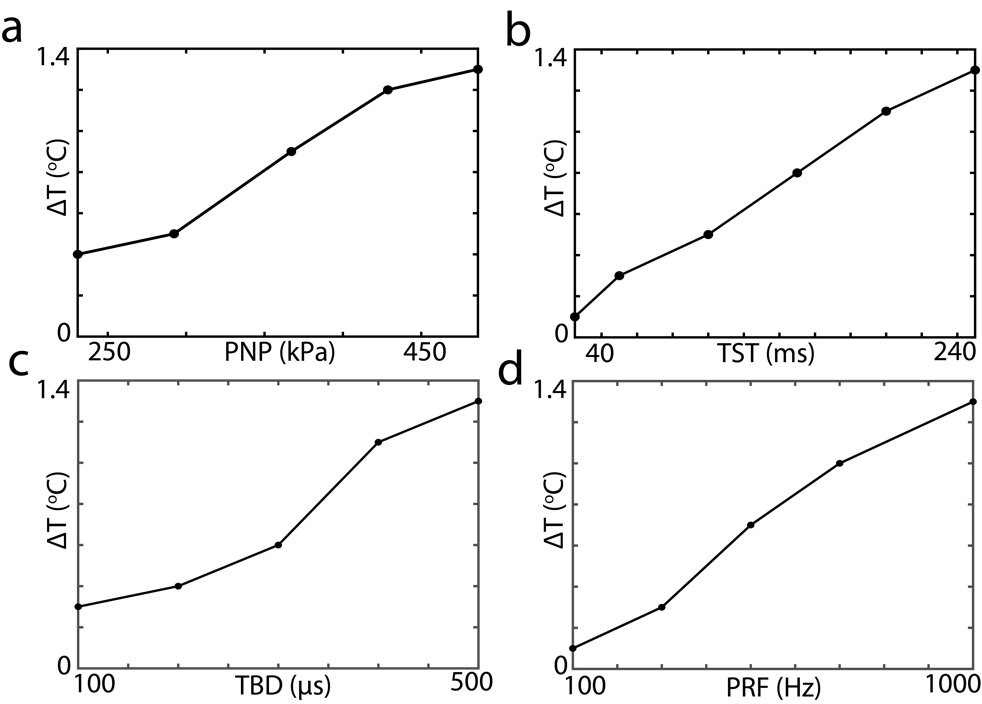

Figure S2: Peak temperature changes with TUT cranial window ultrasound stimulation w.r.t a) peak negative pressure (PNP), b) total stimulation time (TST), c) total burst duration (TBD), and d) pulse repetition frequency (PRF). The parameter space is listed in Table S1.

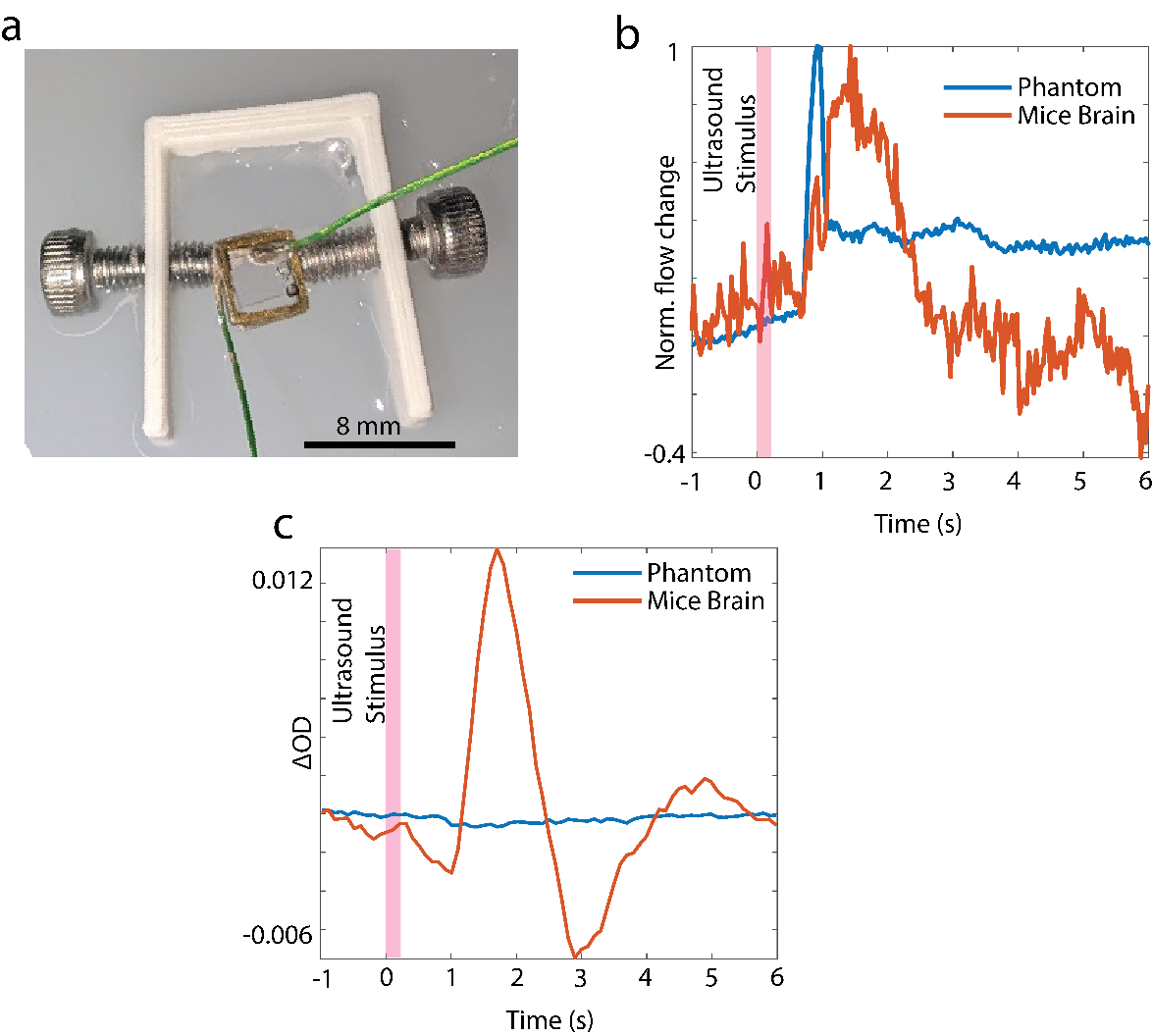

Figure S3: a) TUT embedded on a 1% v/v intralipid phantom. b) The normalized %flow change for phantom (blue) and mice brain (red). The traces are plotted without any moving average. c) shows the optical density (OD) changes of phantom (blue) and mice brain (red).

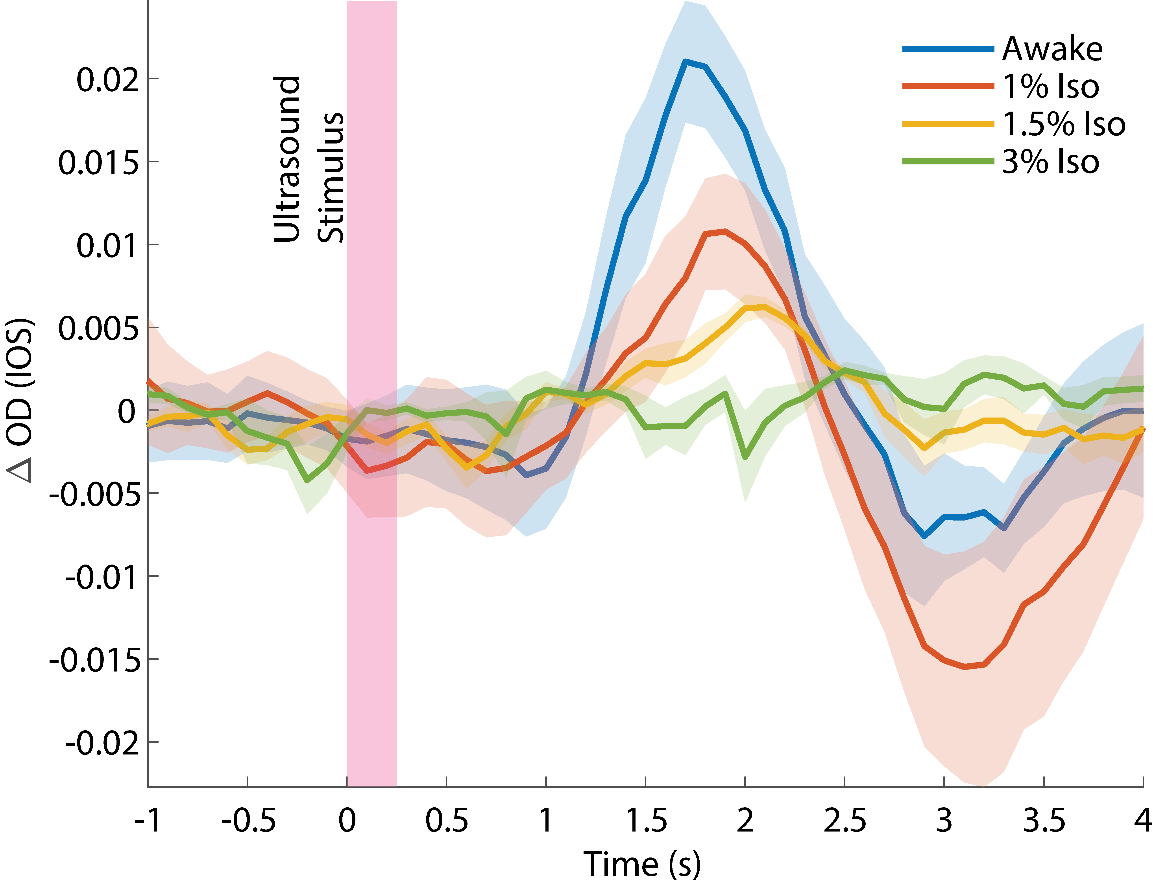

Figure S4: The effects of isoflurane on ultrasound stimulation hemodynamic responses in comparison to awake mice with isoflurane levels varying from 1% to 3% in 1 L/min medical oxygen.

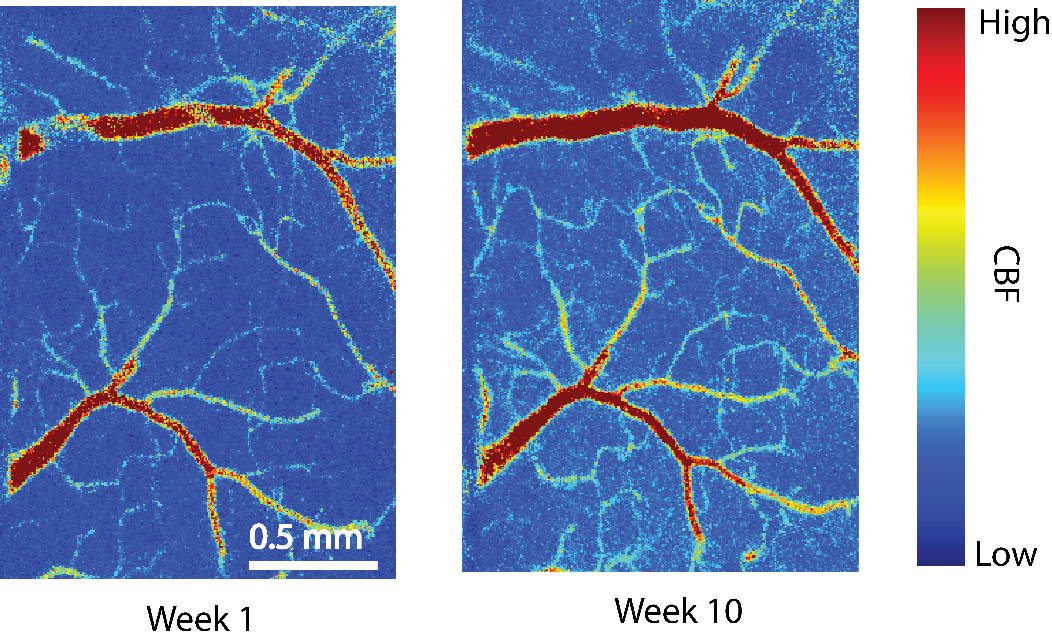

Figure S5: Laser speckle contrast imaging of mice brain through the TUT cranial window over 10 weeks indicating TUT cranial window’s viability for longitudinal imaging.

**Caption for supplementary video 1:**

The video shows the CBF responses recorded through the TUT cranial window in response to ultrasound stimulation at time=0, PRF: 1000 Hz, TBD: 500 μs, PNP: 486.2 kPa, TST: 250 ms.

**Caption for supplementary video 2:**

The video shows the ΔOD responses recorded through the TUT cranial window in response to ultrasound stimulation at time=0, PRF: 1000 Hz, TBD: 500 μs, PNP: 486.2 kPa, TST: 250 ms.

**Table S1: Parameters for ultrasound stimulation used in our study**

| **PNP (kPa)** | **PRF (Hz)** | **TBD (**$\boldsymbol{\mu s}$ | **TST (ms)** | **ISPPA (W/cm^2^)** | $\mathbf{MI}$ |
| --- | --- | --- | --- | --- | --- |
| 486.2 | 1000 | 100 | 250 | 4.9 | 0.121 |
| 486.2 | 1000 | 200 | 250 |  |  |
| 486.2 | 1000 | 300 | 250 |  |  |
| 486.2 | 1000 | 400 | 250 |  |  |
| 486.2 | 1000 | 500 | 250 |  |  |
| 486.2 | 100 | 500 | 250 |  |  |
| 486.2 | 300 | 500 | 250 |  |  |
| 486.2 | 500 | 500 | 250 |  |  |
| 486.2 | 700 | 500 | 250 |  |  |
| 486.2 | 1000 | 500 | 25 |  |  |
| 486.2 | 1000 | 500 | 50 |  |  |
| 486.2 | 1000 | 500 | 100 |  |  |
| 486.2 | 1000 | 500 | 150 |  |  |
| 486.2 | 1000 | 500 | 200 |  |  |
| 230.8 | 1000 | 500 | 250 | 1.3 | 0.062 |
| 292.3 | 1000 | 500 | 250 | 1.9 | 0.076 |
| 367.1 | 1000 | 500 | 250 | 2.8 | 0.092 |
| 428.7 | 1000 | 500 | 250 | 3.9 | 0.108 |

**Table S2: Studies with ultrasound neuromodulation elicited hemodynamic changes in rodents**

| Reference | Transducer | Ultrasound stimulation spot size | Rodent Window | Rodent State | Imaging Technique | Maximum ISPPA  and  Stimulation Time |
| --- | --- | --- | --- | --- | --- | --- |
| [6] | V301-SU, Olympus, USA, f=500 kHz | 6.6 mm x 5.7 mm (Lateral) | Craniotomy surgery  followed by ultrasound stimulation | Anesthesia | Laser speckle contrast imaging | 1.1 W/cm^2^ and 400 ms |
| [7] | H-215; Sonic Concepts, f=4 MHz | 0.5 mm (Lateral)  1.9 mm (Axial) | Craniotomy surgery  followed by ultrasound stimulation | Anesthesia | Functional ultrasound imaging | 360 W/cm^2^ and  10 sec |
| [8] | V323-SU, Olympus, USA  f=2.25 MHz | 6.6 mm (Lateral) | Glass cranial window with craniotomy | Awake | Intrinsic optical signal imaging and Fluorescence Imaging | 6.75 W/cm^2^  and  400 ms |
| [9] | V301-SU, Olympus Corp., Japan, f=425 kHz | 3mm (Lateral)  7.5 mm (Axial) | Intact skull with clear metabond and coverslip | Awake | Intrinsic optical signal imaging | 1.84 W/cm^2^ and  (10 x 200 ms bursts in 5 sec) |
| [10] | PA44LE, Thorlabs  F=623 kHz | 2 mm (Lateral)  5 mm (Axial) | Fiber implantation | Freely moving | Fiber Photometry | 7.4 W/cm^2^  and  10 sec |
| [11] | V301-SU, Olympus, USA, f=500 kHz | 6.9 mm (Lateral) | Glass cranial window with craniotomy | Awake and Anesthesia | Intrinsic optical signal imaging | 27 W/cm^2^ and  400 ms |
| Our study | In house fabricated lithium niobate TUT, f=12 MHz | 3 mm  (Lateral) | Thinned skull TUT cranial window | Awake | Laser speckle contrast imaging  And Intrinsic optical signal imaging | 4.9 W/cm^2^  and  250 ms |
